# Macrophage-based assays for the in vitro testing of the anti-inflammatory activity of mesenchymal stem cell-based products

**DOI:** 10.64898/2026.03.04.709479

**Authors:** Andrea Exnerová, Sabina Seidlová, Věra Daňková, Vojtěch Pavlík, Kristina Nešporová

## Abstract

Therapies based on mesenchymal stromal cells (MSCs) have high potential in the field of regenerative medicine due mainly to their immunomodulatory properties. However, their clinical translation is hampered by a lack of sufficiently standardised potency tests. Since macrophages comprise key mediators of the effects of MSCs, macrophage-based assays potentially provide a relevant in vitro tool for the evaluation of the activity of MSC products. This study involved the coculturing of canine adipose-derived mesenchymal stem cells (ASCs) with macrophages derived from human THP-1 and U937 monocyte cell lines, murine RAW264.7 macrophages and primary human macrophages. The M2 polarisation was assessed following stimulation with IL-4/IL-13. The mRNA expression of the pro- and anti-inflammatory markers was analysed applying qPCR. The ASC secretome acted to reduce the pro-inflammatory mRNA expression across all the macrophage models, albeit with a certain degree of model–dependent variability. Only the U937 macrophages responded consistently to the M2-polarising stimuli, while the RAW264.7 cells provided practical advantages in terms of routine screening. The results thus provided support for the application of macrophage-based potency assays as a suitable platform for the testing of MSC products; the U937 cells were found to be particularly suitable for the study of polarisation and the RAW264.7 cells for standardised screening.

## Introduction

MSCs comprise adult multipotent stromal stem cells that can be isolated from a wide range of tissues, including bone marrow, adipose tissue and dental pulp. MSCs have demonstrated their therapeutic potential in terms of a broad range of human and veterinary diseases, including osteoarthritis, chronic wounds, graft-versus-host disease (GvHD), Crohn’s disease and neurological disorders (1,2). Extensive preclinical studies have further demonstrated that MSCs possess immunomodulatory and pro-regenerative properties (3). ASCs comprise a subset of MSCs, and they exhibit regenerative properties that are broadly comparable to those of MSCs obtained from other sources.

Translating MSC therapy into clinical practice requires the application of standardised production and quality control approaches. The International Society for Cell & Gene Therapy sets out minimum criteria for the characterisation of MSCs, including adherence to tissue culture plastic, the expression of CD105, CD73 and CD90, a lack of haematopoietic markers and trilineage differentiation potential (5). The conventional clinical quality control approach primarily addresses identity, viability and sterility; however, donor-to-donor and batch-to-batch variability necessitate the conducting of additional potency tests (6–8). Potency assays should be indication-specific and should, ideally, correlate with therapeutic efficacy.

The potency of MSCs can be assessed directly applying gene, protein or surface marker analyses, or indirectly via functional assays that measure the impacts of MSCs on target cells (6,9). Mixed lymphocyte reactions (MLRs) assess the interactions of MSCs with T cells and are relevant for diseases that are driven by adaptive immunity, such as GvHD. However, with respect to conditions such as osteoarthritis or chronic wounds, macrophages often comprise the key dysregulated immune cells, and MSCs are thought to primarily modulate macrophage phenotypes (10–12). Consequently, the development of macrophage-based potency assays is steadily accelerating (13).

This study presents macrophage-based assays that make use of immune cell lines to assess ASC immunomodulatory potency via the quantification of the expression of pro-inflammatory and pro-regenerative mRNA using qPCR. We employed canine ASCs aimed at supporting potential translation to veterinary clinical trials.

## Methods

### Ethical approval statement

Adipose tissue (approximately 3 g) was obtained in the form of a surplus material during routine, medically indicated veterinary procedures in client-owned dogs. No procedures were performed for research purposes. According to Czech legislation (Act No. 246/1992 131 Coll., on the Protection of Animals Against Cruelty), animals that are subjected to diagnostic or therapeutic interventions are not considered to be experimental animals; therefore, no approval was required from the Institutional Animal Care and Use Committee. All the veterinary procedures were performed by licensed veterinarians, and the owners provided their informed consent for the use of any surplus tissue for research purposes. All the experiments were performed in vitro using isolated cells.

The primary human macrophages were derived from peripheral blood monocytes obtained from healthy donors following their informed consent and in accordance with the relevant institutional and national ethical requirements.

### Isolation and culture of the canine ASCs

Fat tissue samples were obtained during the conducting of routine ovariectomies in female dogs. A portion of the surplus visceral fat (approximately 3 g) was washed in 9.8 ml of physiological saline supplemented with 200 µl of penicillin-streptomycin (Diagnovum), minced and digested with 2 mg/ml of type I collagenase (Gibco) dissolved in αMEM (Capricorn Scientific) containing 2% penicillin/streptomycin at 37°C, 200 rpm for 60 min. The suspension was filtered through a 40 µm mesh, diluted with 10 ml of αMEM and centrifuged at 300 g for 5 min. The pellets were washed with physiological saline, resuspended in αMEM containing 10% FBS (Biosera), 1% stable L-glutamine (Carl Roth) and 1% penicillin/streptomycin, and seeded in 150 cm^2^ flasks (total volume of 20 ml). The cells were cultured at 37°C in 5% CO_2_, the medium was changed twice per week and the cells were passaged at 70%–80% confluence.

### U937 and THP-1 cell culture and macrophage differentiation

The U937 (human acute monocytic leukaemia cell line, Merck, cat. # 85011440-1VL) and THP-1 (human acute monocytic leukaemia cell line, Merck, cat. # 88081201-1VL) cells were cultured in RPMI 1640 medium (Biosera) supplemented with 10% FBS, 2 mM L-glutamine, 1% penicillin/streptomycin and 1 mM sodium pyruvate (Merck); the THP-1 medium further contained 4.5 g/l D-glucose (Thermo Fisher) and 0.05 mM β-mercaptoethanol (Merck). The U937 cells were seeded at 5 × 10^5^ cells/ml and THP-1 cells at 2.5 × 10^5^ cells/ml. For differentiation purposes, both the U937 (1 × 10^6^ cells/well) and THP-1 (5 × 10^5^ cells/well) cells were treated with 50 ng/ml PMA (Merck) for 96 h and 24 h, respectively, then cultured in PMA-free medium for 24 h prior to the conducting of the coculture experiments.

### RAW264.7 cell culture

The RAW264.7 cells (murine tumour macrophage-like cell line, ATCC, cat. # TIB-71) were maintained in DMEM (Biosera) supplemented with 10% FBS, 2 mM L-glutamine and 1% penicillin/streptomycin. For experimental purposes, 5 × 10^5^ cells were seeded per well in 24-well plates and used the following day.

### Generation of primary human macrophages

The peripheral blood monocytes were cultured in RPMI 1640 medium supplemented as described above for the THP-1 cells and differentiated for 6 days with 50 ng/ml M-CSF. The medium was replaced every 2–3 days.

### M2 macrophage activation

The macrophages (U937, THP-1, RAW264.7) were stimulated using IL-4 and IL-13 (25 ng/ml each) for 24 and 48 h. Following incubation, the cells were lysed for the RNA isolation and qPCR analyses. The experiments were performed in triplicate.

### Coculture of ASCs and macrophages

The ASCs were cocultured with macrophages using 0.4 µm pore-size PET inserts so as to allow for paracrine signalling without direct contact with the cells. The macrophages were stimulated with LPS (from *Pseudomonas aeruginosa*) at 100 ng/ml for the RAW264.7 and U937-derived macrophages, and at 1,000 ng/ml for the THP-1-derived macrophages. These concentrations were selected based on commonly applied conditions as reported in the literature and were refined for the preliminary optimisation experiments so as to attain the robust and reproducible induction of inflammatory markers in each of the models.

The ASC-to-macrophage ratios were: RAW264.7—1:1, 1:2 and 1:5; U937—1:2; THP-1—1:1. The ratios were adjusted in order to account for differences in the cell sizes and plating requirements across the models and to avoid the overconfluence or stress of the ASCs on the transwell inserts. The cocultures were performed for 24 and 48 h; these time points were chosen aimed at capturing the early and intermediate transcriptional responses while maintaining feasibility for routine screening purposes. Following incubation, the macrophages were harvested for RNA isolation. Each of the conditions was tested in the form of ≥3 independent biological replicates, as indicated in the figure legends.

### Quantitative PCR (qPCR)

The total RNA was extracted using an RNeasy Mini Kit (Qiagen). The cDNA was synthesised, and TaqMan qPCR was performed (QuantStudio 3, Thermo Fisher Scientific). The target mRNAs for the M2 activation comprised *TGFβ1* (Hs00998133_m1), *IL10* (Hs00961622_m1), *CCL22* (Hs01574247_m1), *CD209* (Hs01588349_m1) and *IL1RA* (Hs00893626_m1); and for inflammation comprised *TNFα* (Hs00174128_m1), *IL1β* (Hs01555410_m1) and *PTGS2* (Hs00153133_m1). The expression was normalised to *GAPDH* and calculated applying the 2^– ΔΔCt method.

### Statistical analysis

The analysis of the data was conducted employing GraphPad Prism software. One-sample t-tests were applied when the values were compared to defined reference values (set at 100%). Paired t-tests were employed to compare differing ASC-to-macrophage ratios when determined in the individual experiments. The data were presented as the mean ± SD from independent biological replicates (n is stated in each of the figure legends). The macrophages are denoted as MΦ in the figures. Concerning the LPS experiments, the significance of MΦ vs MΦ + LPS is indicated by #, whereas the significance of MΦ + LPS vs MΦ + LPS + ASC is indicated by *. With regard to the IL-4/IL-13 stimulation, the significance of MΦ vs MΦ + IL-4 + IL-13 is indicated by #, and the significance of MΦ + IL-4 + IL-13 vs MΦ + IL-4 + IL-13 + ASC (when applicable) is indicated by *. The significance levels are denoted as #/* p ≤ 0.05, ##/** p ≤ 0.01, ###/*** p ≤ 0.001, ####/**** p ≤ 0.0001; ns, not significant.

## Results

The U937-derived macrophages, THP-1-derived macrophages, RAW264.7 macrophages and primary human macrophages were stimulated with LPS in the presence or absence of the ASC coculture using transwell inserts. The mRNA expression of the pro-inflammatory markers was analysed after 24 and 48 h. The ASC coculture acted to reduce the LPS-induced expression of pro-inflammatory genes across all the tested macrophage models, although the magnitude and temporal patterns of inhibition varied between the cell lines. For example, the modulation of the TNFα expression at 48 h differed between the U937 and THP-1 macrophages when compared to the RAW264.7 cells (Fig. 1A–I). The ASC secretome further suppressed the pro-inflammatory mRNA expression in the U937 macrophages when applied after a short period (3 h) of LPS pre-activation, thus indicating activity in the already-activated macrophages (Fig. S1A–C). Comparable anti-inflammatory effects were confirmed in the primary human macrophages derived from the peripheral blood monocytes (Fig. 2).

**Figure 1.**
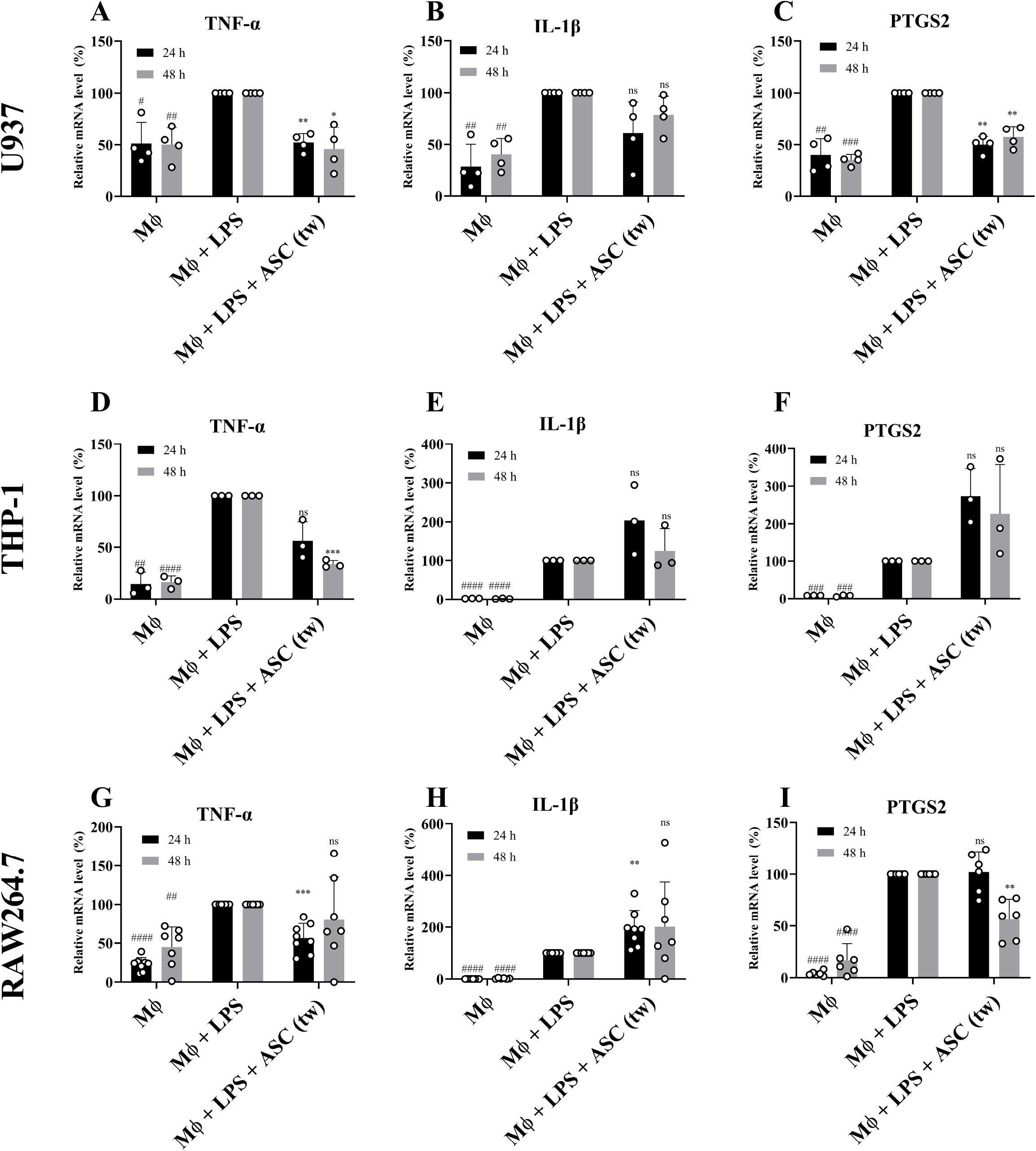
The anti-inflammatory effect of ASC secretome seeded in transwells (tw) in various LPS-stimulated macrophage models. A-C: the mRNA expression of inflammatory markers in the U937 macrophages. D-F: the mRNA expression of inflammatory markers in the THP-1 macrophages. G-I: the mRNA expression of inflammatory markers in the RAW264.7 macrophages. A-F: The data are presented as the mean relative mRNA expression ± standard deviation from four independent experiments. G-I: The data are presented as the mean relative mRNA expression ± standard deviation from seven independent experiments. The statistical analysis was performed applying the one-sample t-test (comparison with the reference value of 100%). The statistical significance of MΦ versus MΦ + LPS is displayed as “#” and the statistical significance of MΦ + LPS versus MΦ + LPS + ASC is displayed as “*”. The significance is indicated as *p ≤ 0.05, **p ≤ 0.01, ***p ≤ 0.001, ****p ≤ 0.0001; ns, not significant (# is used analogically).

**Figure 2.**
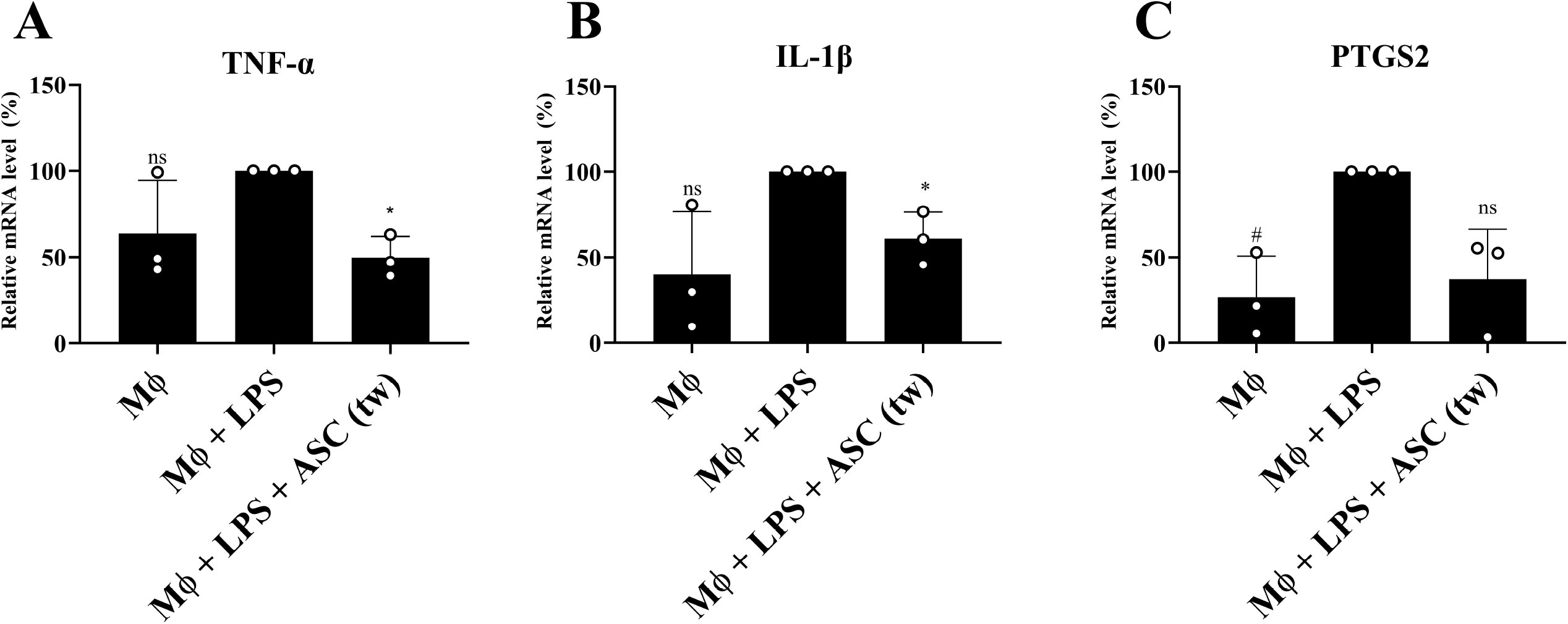
The anti-inflammatory effect of the ASC secretome in human peripheral-blood-monocyte-derived macrophages. The data are presented as the mean relative mRNA expression ± standard deviation from three independent experiments. The statistical analysis was performed applying the one-sample t-test (comparison with the reference value of 100%). The statistical significance of MΦ versus MΦ + LPS is displayed as “#” and the statistical significance of MΦ + LPS versus MΦ + LPS + ASC is displayed as “*”. The significance is indicated as *p ≤ 0.05, **p ≤ 0.01, ***p ≤ 0.001, ****p ≤ 0.0001; ns, not significant (# is used analogically).

Given their adherent growth and lack of differentiation requirements, the RAW264.7 cells were selected for the further evaluation of the ASC dose dependency. Increasing the ASC-to-macrophage ratios resulted in the dose-dependent modulation of the pro-inflammatory mRNA expression. A statistically significant difference was observed between the 1:1 and 1:2 ratios, whereas no significant difference was observed between the 1:2 and 1:5 ratios (Fig. 3).

**Figure 3.**
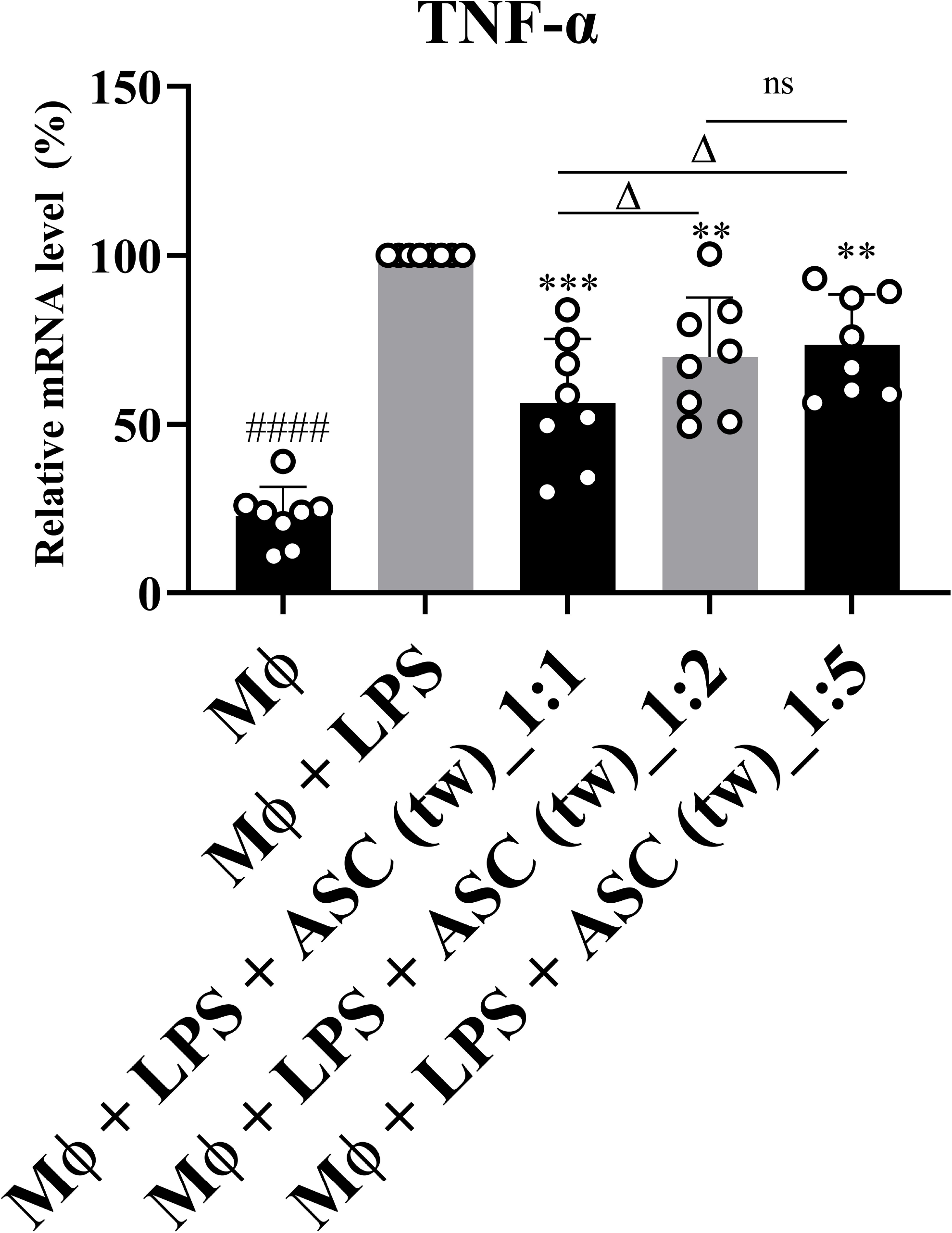
The concentration dependency of the anti-inflammatory effect of the MSC secretome. The data are presented as the mean relative mRNA expression ± the standard deviation from eight independent experiments. The statistical analysis was performed applying the one-sample t-test (comparison with the reference value of 100%). The statistical significance of MΦ versus MΦ + LPS is displayed as “#” and the statistical significance of MΦ + LPS versus MΦ + LPS + ASC is displayed as “*”. The significance is indicated as *p ≤ 0.05, **p ≤ 0.01, ***p ≤ 0.001, ****p ≤ 0.0001; ns, not significant (# is used analogically). The paired t-test was used for the statistical evaluation of the differing ASC (tw) ratios; Δ p≤ 0.05, ns – not significant.

Aimed at evaluating whether the macrophage models were suitable for the assessment of regenerative polarisation, they were stimulated with IL-4 and IL-13, followed by the analysis of the M2-associated gene expression. Only the U937-derived macrophages exhibited the consistent upregulation of IL-10 and TGFβ1 following IL-4/IL-13 stimulation (Fig. 4). In contrast, the RAW264.7 and THP-1 macrophages did not exhibit the reproducible transcriptional induction of these markers under the test conditions. The further analysis of the U937 macrophages demonstrated that the ASC secretome modestly potentiated IL-4/IL-13-induced M2-associated mRNA expression, including IL1RA, CD209 and CCL22 (Fig. 5). In addition, the ASC secretome further reduced the expression of selected pro-inflammatory markers in the IL-4/IL-13-polarised macrophages (Fig. S2). However, the ASC secretome alone was insufficient in terms of inducing M2-associated mRNA expression in the short-term LPS-preactivated U937 macrophages (Fig. S1D–H).

**Figure 4.**
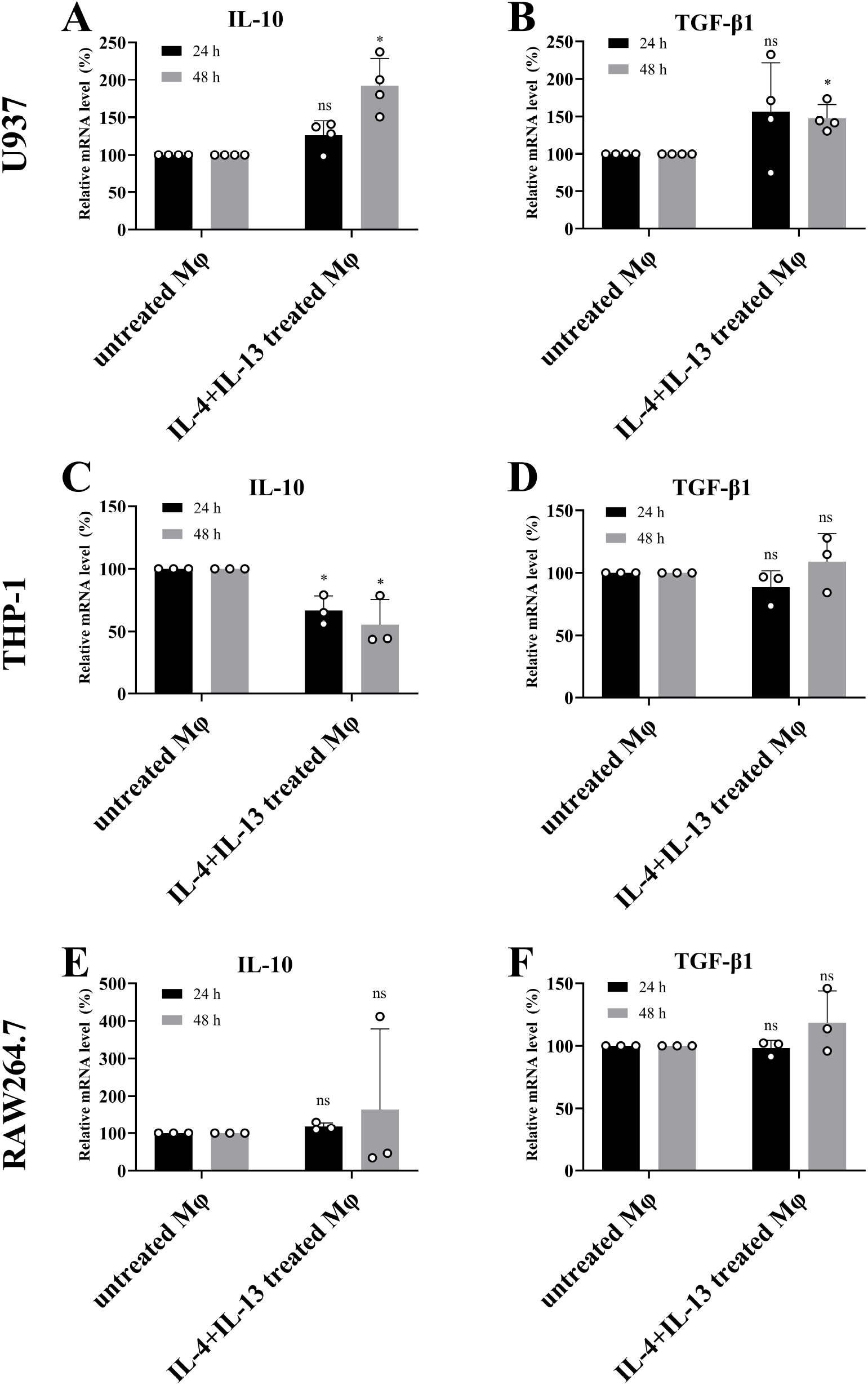
The activation of the various macrophage cell lines in the M2 phenotype. A, B: The data are presented as the mean relative mRNA expression ± the standard deviation from four independent experiments. C-F: The data are presented as the mean relative mRNA expression ± the standard deviation from three independent experiments. The statistical analysis was performed applying the one-sample t-test (comparison with the reference value of 100%). The statistical significance of MΦ versus MΦ + IL-4 + IL-13 is displayed as “**\***”. The significance is indicated as *p ≤ 0.05, **p ≤ 0.01, ***p ≤ 0.001, ****p ≤ 0.0001; ns, not significant.

**Figure 5.**
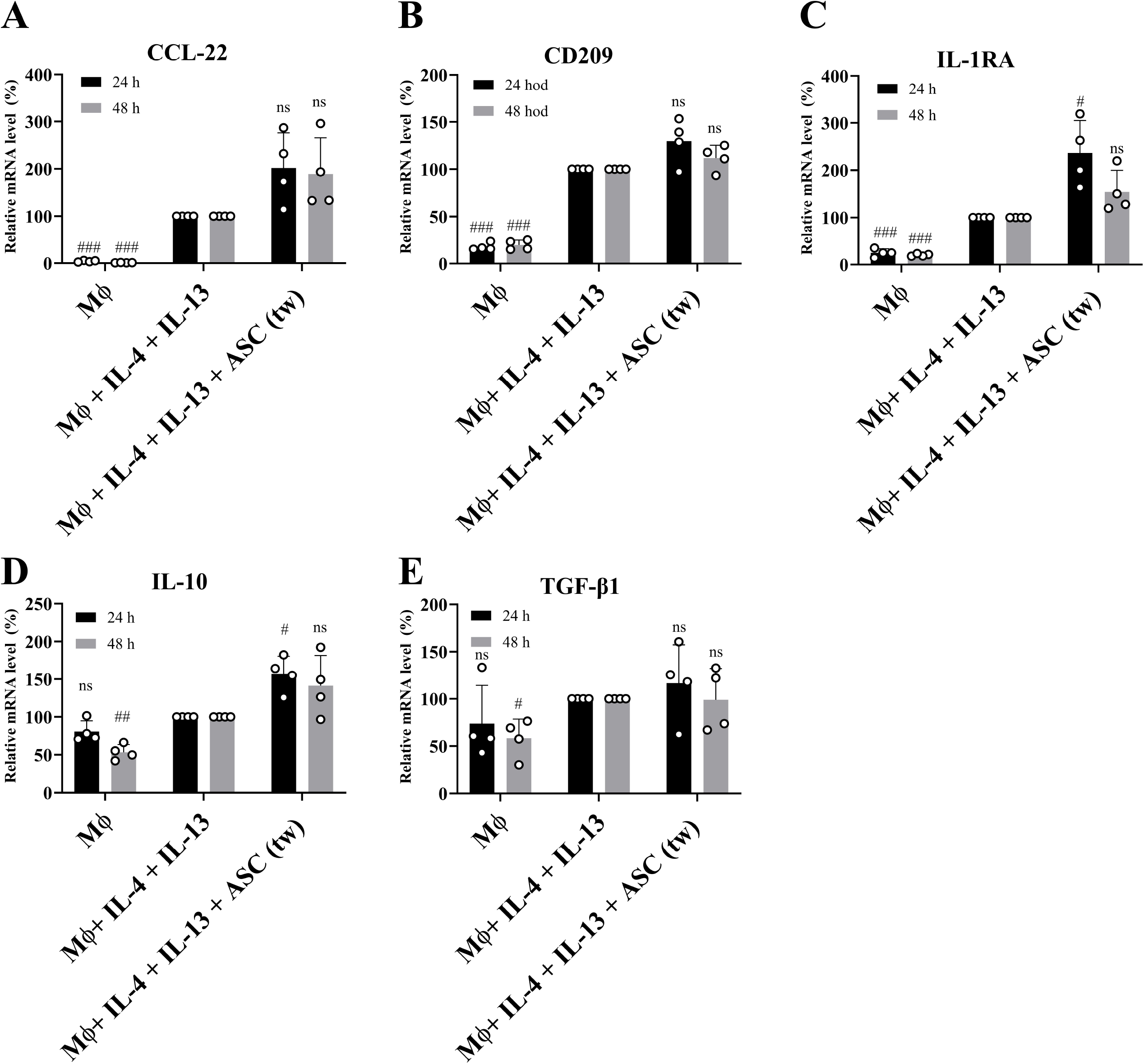
The effect of the ASC secretome on the M2 polarisation of U937. A-E: the mRNA expression of the various M2 markers. The data are presented as the mean relative mRNA expression ± the standard deviation from four independent experiments. The statistical analysis was performed applying the one-sample t-test (comparison with the reference value of 100%). The statistical significance of MΦ versus MΦ + IL-4 + IL-13 is displayed as “#” and the statistical significance of MΦ + IL-4 + IL-13 versus MΦ + IL-4 + IL-13 + ASC is displayed as “*”. The significance is indicated as *p ≤ 0.05, **p ≤ 0.01, ***p ≤ 0.001, ****p ≤ 0.0001; ns, not significant (# is used analogically).

## Discussion

Although mesenchymal stem cell (MSC)-based products have been subjected to detailed investigation in the field of regenerative medicine in recent years, their clinical translation remains limited due to suboptimal dosing and timing and a lack of standardised manufacturing and quality control approaches (2,4,14–16). While MSCs exhibit well-established immunomodulatory and anti-inflammatory effects, their successful clinical implementation depends on the optimisation of robust potency assays, the dosing strategy and consistent product characterisation (17,18).

Potency testing has become a key component in terms of the evaluation of MSC products. Traditional T-cell proliferation assays are beset with challenges surrounding both their variability and limited reproducibility (9). Recent evidence indicates that macrophages (rather than lymphocytes) comprise the primary immune targets of MSC products and their secretomes (12). Consequently, macrophage-based assays are gaining relevance, especially with regard to conditions such as wound healing and osteoarthritis, in which macrophages orchestrate both the inflammatory and reparative processes (10,19,20).

THP-1, U937 and RAW264.7 rank amongst the most commonly applied macrophage models. THP-1 and U937 comprise human monocyte-derived cell lines with distinct differentiation kinetics, while RAW264.7 cells comprise an immortalised murine macrophage line with a relatively pro-inflammatory, M1-like baseline phenotype. M2 polarisation, which is typically induced by IL-4 and IL-13, is used to assess the pro-regenerative potential; however, the degree of efficiency of polarisation varies substantially across cell lines (21–24). In line with this observation, the polarisation responses differed markedly across the tested models, as previously reported in studies that determined substantial heterogeneity between commonly used macrophage lines. THP-1 and U937 differ in terms of their differentiation protocols and basal activation states, which potentially significantly condition cytokine-driven transcriptional programs. RAW264.7 cells frequently display a relatively high basal inflammatory tone, which may act to attenuate responses driven by IL-4/IL-13 under standardised conditions. Together, these observations reinforce the fact that macrophage models are not functionally interchangeable and should be benchmarked and validated prior to their use for standardised potency testing purposes.

Although this study confirmed the anti-inflammatory role of the ASC secretome in various macrophage models, it also highlighted certain line-specific responses. IL-1β inhibition was detected only for the U937 macrophages, which possibly reflected differences in the LPS sensitivity or higher intrinsic pro-inflammatory bias in the other lines. Similarly, the PTGS2 expression in the THP-1 cells was observed not to be suppressed, most likely due to stronger LPS-induced activation. The primary human macrophages confirmed a concentration-dependent anti-inflammatory effect, thus supporting the potential use of this setup for standardised potency testing purposes.

Only the U937 macrophages responded to IL-4 and IL-13 stimulation, with the upregulation of IL-10 and TGF-β1, thus suggesting that this line best represents M2-like macrophages. Although protein-level analyses and surface marker profiling have the potential to refine this model, qPCR-based assays provide a practical and scalable framework for preclinical evaluation purposes. Our findings indicate that the ASC secretome predominantly exerts anti-inflammatory rather than M2-polarising effects, which suggests that the reported M2 induction may be a secondary consequence of the suppression of inflammation. The main aim of this study was to compare various cell lines that are suitable for the routine biological testing of MSC-based products. It was not our intention to demonstrate pronounced phenotypic shifts, but rather to evaluate the sensitivity and consistency of various macrophage models as readouts for the activity of MSC secretomes. In this context, the modest yet reproducible transcriptional changes in the key immunoregulatory markers are informative in terms of the assessment of the comparative potency rather than for mechanistic interpretation purposes.

The exclusive reliance on transcriptional readouts represents one of the limitations of the study. Although qPCR-based assays provide a high degree of sensitivity and are well-suited to scalable screening workflows, mRNA levels do not necessarily reflect the protein expression or biological function. Additional validation at the protein level (e.g. the quantification of secreted cytokines or the intracellular protein expression) and functional assays that assess the behaviour of macrophages are required in order to be able to fully characterise polarisation states. Moreover, intervention-based approaches such as neutralisation or blocking experiments are required so as to identify the specific mediators within the ASC secretome that are responsible for the observed effects. Since our study was designed to provide a comparative evaluation of selected macrophage models rather than a mechanistic investigation, these aspects were beyond the scope of the research and should be addressed in future studies.

From the translational perspective, the proposed macrophage-based platform represents progress towards the development of functionally oriented, GMP-compliant assays for MSC standardisation and batch release purposes. Incorporating a biologically meaningful readout that reflects the immunomodulatory activity of the MSC secretome would complement the established identity, purity and viability criteria and provide support for a more comprehensive evaluation of product consistency. Such an approach would be particularly valuable given the well-documented donor-to-donor and manufacturing-associated variability of MSC preparations.

Nevertheless, it is necessary to acknowledge several limitations and practical considerations. The intrinsic biological variability of both MSC products and responder macrophage models potentially influences the performance of assays and, therefore, requires a careful control approach. The establishment of assay robustness and reproducibility, as well as clearly defined acceptance criteria, represent essential steps towards validation. In addition, the fact that compliance with regulatory expectations concerning potency assays in the GMP framework— including the standardisation and qualification of assays and the definition of release specifications—is critical prior to routine implementation in the clinical manufacturing environment should be carefully considered.

Finally, the use of canine ASCs with human and murine macrophages illustrates the potential applicability of this assay platform for the testing of xenogeneic MSC-based therapeutics across species. This cross-species approach was intentionally adopted as a pragmatic, exploratory strategy for the evaluation of the robustness of assays across distinct responder systems. While such models may be suitable for early-stage screening or veterinary applications, caution is required when extrapolating these findings to human clinical settings. Species-specific differences in cytokine networks, receptor-ligand affinity and downstream signalling pathways potentially influence both the magnitude and the qualitative nature of the observed responses. Therefore, although the results presented in this study demonstrate functional compatibility at the transcriptional level, species-matched validation is required before translating this platform into a human GMP clinical manufacturing framework.

## Conclusion

MSC-based therapies are advancing towards clinical applications, which underscores the need for standardised and functionally meaningful testing aimed at ensuring reproducibility across studies and research facilities. Our findings provide foundational guidance for the selection and benchmarking of macrophage models that support standardisation. Facilities that research such assays must employ thorough validation approaches (including assay qualification and the definition of acceptance criteria) in accordance with applicable regional regulatory requirements.

## Availability of the data and materials

The data generated in the study are available at 10.6084/m9.figshare.31484923.

## Authors’ contributions

AE - investigation, methodology, formal analysis, writing – original draft, SS – investigation, formal analysis, VD – writing – review and editing, VP – writing – review and editing, KN - conceptualisation, supervision

## Ethics approval and consent to participate

The adipose tissue used for the isolation of the canine MSCs was obtained in the form of surgical waste during planned procedures, with the informed consent of the owners.

## Patient consent for publication

Not applicable

## Acknowledgments

The authors would like to thank the technical staff of the Cell physiology group for their invaluable assistance with the preparation of the samples and collection of the data. The authors sincerely acknowledge Doc. Vladimir Velebny for his leadership and for enabling the conducting of the research.

## Supplementary data

**Figure S1.**
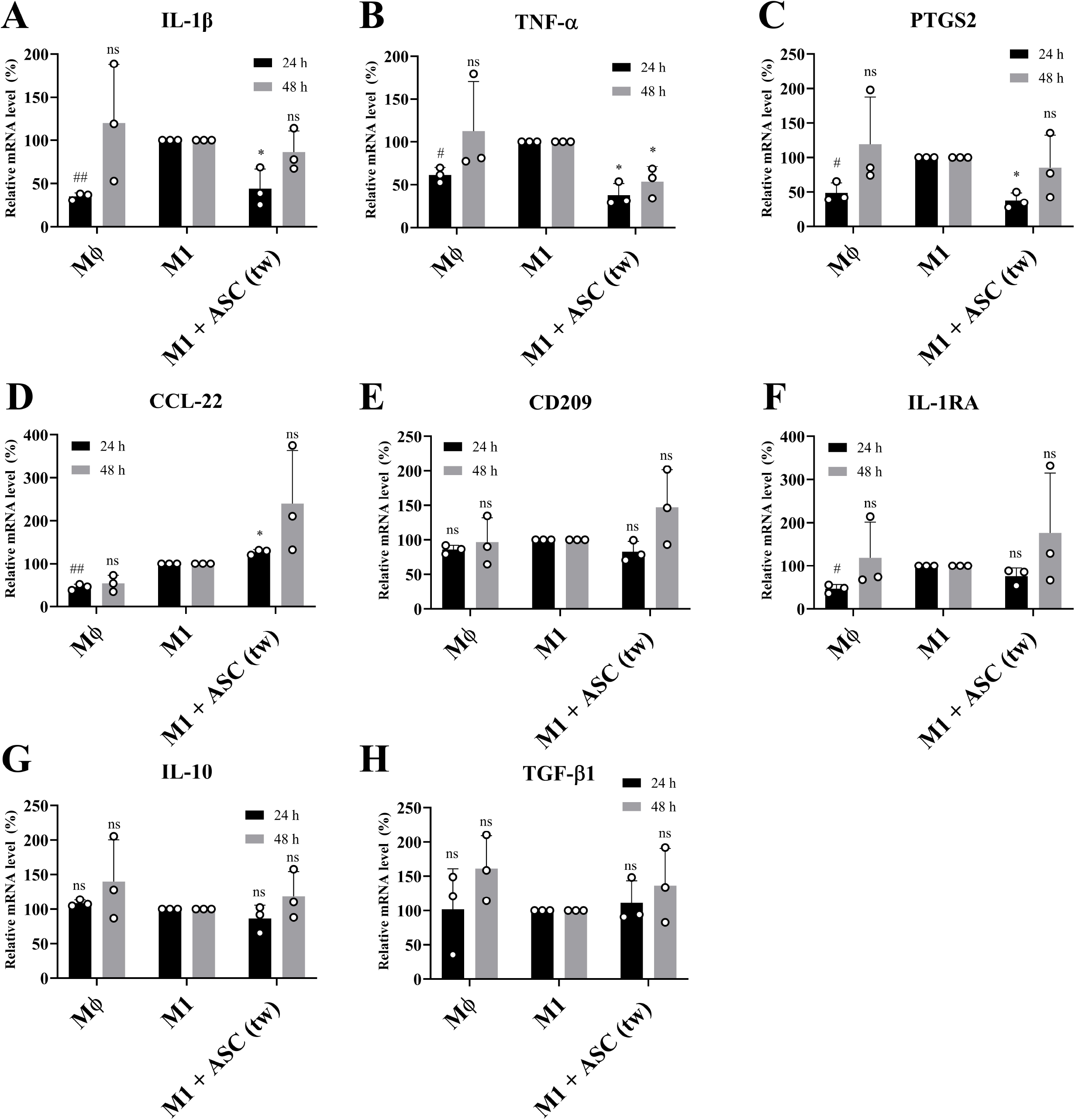
the anti-inflammatory effect of the ASC secretome on the LPS-pre-activated U937 macrophages. The U937 cells were briefly (3 h) pre-activated with LPS prior to the seeding of the ASC. A-C: The effect of the ASC secretome on the mRNA expression of the M1 markers. D-H: The effect of the ASC secretome on the mRNA expression of the M2 markers. The data are presented as the mean relative mRNA expression ± the standard deviation from three independent experiments. The statistical analysis was performed applying the one-sample t-test (comparison with the reference value of 100%). The statistical significance of MΦ versus MΦ + IL-4 + IL-13 is displayed as “#” and the statistical significance of MΦ + IL-4 + IL-13 versus MΦ + IL-4 + IL-13 + ASC is displayed as “*”. The significance is indicated as *p ≤ 0.05, **p ≤ 0.01, ***p ≤ 0.001, ****p ≤ 0.0001; ns, not significant (# is used analogically).

**Figure S 2.**
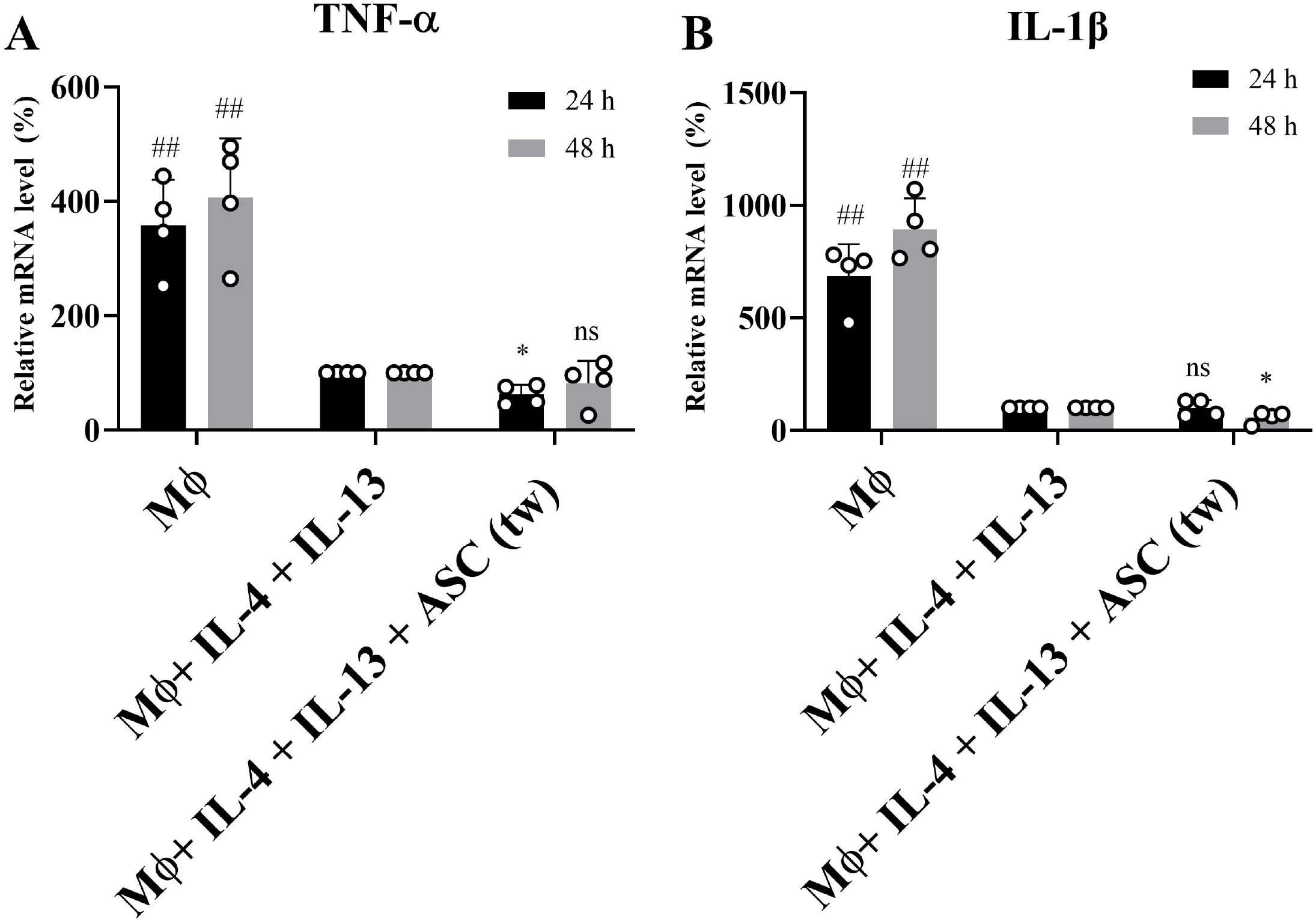
The effect of the ASC secretome on the mRNA expression of the M1 markers in the U937 M2 macrophage model. The data are presented as the mean relative mRNA expression ± the standard deviation from four independent experiments. The statistical analysis was performed applying the one-sample t-test (comparison with the reference value of 100%). The statistical significance of MΦ versus MΦ + IL-4 + IL-13 is displayed as “#” and the statistical significance of MΦ + IL-4 + IL-13 versus MΦ + IL-4 + IL-13 + ASC is displayed as “*”. The significance is indicated as *p ≤ 0.05, **p ≤ 0.01, ***p ≤ 0.001, ****p ≤ 0.0001; ns, not significant (# is used analogically).

